# The Oft-Overlooked Massively Parallel Reporter Assay: Where, When, and Which Psychiatric Genetic Variants are Functional?

**DOI:** 10.1101/2020.02.02.931337

**Authors:** Bernard Mulvey, Tomas Lagunas, Joseph D. Dougherty

**Author notes:** Contact: Dr. Joseph Dougherty, Dougherty Lab, Department of Genetics, 660 S. Euclid Ave, Campus Box 8232, St. Louis, MO 63110-1093, E.

## Abstract

Neuropsychiatric phenotypes have been long known to be influenced by heritable risk factors. The past decade of genetic studies have confirmed this directly, revealing specific common and rare genetic variants enriched in disease cohorts. However, the early hope for these studies—that only a small set of genes would be responsible for a given disorder—proved false. The picture that has emerged is far more complex: a given disorder may be influenced by myriad coding and noncoding variants of small effect size, and/or by rare but severe variants of large effect size, many *de novo*. Noncoding genomic sequences harbor a large portion of these variants, the molecular functions of which cannot usually be inferred from sequence alone. This creates a substantial barrier to understanding the higher-order molecular and biological systems underlying disease risk. Fortunately, a proliferation of genetic technologies—namely, scalable oligonucleotide synthesis, high-throughput RNA sequencing, CRISPR, and CRISPR derivatives—have opened novel avenues to experimentally identify biologically significant variants *en masse*. These advances have yielded an especially versatile technique adaptable to large-scale functional assays of variation in both untranscribed and untranslated regulatory features: Massively Parallel Reporter Assays (MPRAs). MPRAs are powerful molecular genetic tools that can be used to screen tens of thousands of predefined sequences for functional effects in a single experiment. This approach has several ideal features for psychiatric genetics, but remains underutilized in the field to date. To emphasize the opportunities MPRA holds for dissecting psychiatric polygenicity, we review here its applications in the literature, discuss its ability to test several biological variables implicated in psychiatric disorders, illustrate this flexibility with a proof-of-principle, *in vivo* cell-type specific implementation of the assay, and envision future outcomes of applying MPRA to both computational and experimental neurogenetics.

## Introduction

Psychiatric diseases are genetically influenced by common and rare heritable variation, as well as by non-inherited, *de novo* mutations. Liability to disease through common variants (allele frequency ≥ 1%) ranges from 10-25% for psychiatric disorders including major depressive disorder (MDD) (1–3) and schizophrenia (1,4). Rare variants have likewise been shown to confer disease risk, particularly in neurodevelopmental and psychotic (5,6) domains. While loci containing both types of variants have been associated with neuropsychiatric phenotypes, two major hurdles have prevented use of this knowledge to better understand disease mechanisms: 1) a large volume of GWAS associations and rare variant discoveries to test for functionality/causality; and 2) the lack of means to predict a variant’s functional consequences. These hurdles confound the identification of shared features across functional variants that converge on common phenotypes.

### Defining specific causal variant(s): problems of linkage and numbers

The identification of candidate genetic variants with disease associations relies on correlational methodologies. Genome wide association studies (GWAS) identify loci with overrepresented blocks of linked variants, comprising numerous single nucleotide polymorphisms (SNPs). Similarly, family studies (*e.g.*, trio studies) identify many proband-specific (*de novo*) or -enriched (rare, inherited) variants in each patient, though only 1-2 may be causal. Further, while consequences of coding variants (*e.g.*, nonsense mutations) can be reliably predicted, the vast majority of GWAS loci fall outside of transcribed sequences(7). Rare and *de novo* variation, on the other hand, is more broadly distributed across the genome, including un*translated* and translation-regulating sequences, well-illustrated by autism spectrum disorders (ASDs): over 255,106 distinct *de novo* variants (including 142 stop codon gains, 3,402 in untranslated regions (UTRs), and 6,787 upstream gene/promoter variants) were recently identified in 1,902 ASD subjects (8). However, these discovery-oriented approaches are incapable of identifying the minority of associations corresponding to variants with biological/disease-pertinent functions.

### Challenges in predicting a variant’s functional consequences

Predicting whether and how a noncoding variant is functional is a nontrivial enterprise. The majority of these variants, and their linked neighbors, bear indirect indication(s) of transcriptional regulatory function in annotations such as expression quantitative trait locus (eQTL) associations, chromatin accessibility, or histone marks (7,9,10). However, even within a cell or tissue type, such data are often mutually discordant: one study examining six epigenomic datasets in K562 cells found that 49% of functional regulators did not overlap *any* of the six annotation sets, and another 40% only overlapped one of the six (11). Similarly, the Roadmap Epigenomics Consortium (12) inferred cell type-specific regulatory elements using chromatin marks, but only a minority of GWAS SNPs overlap these elements (except in blood) (13). Therefore, epigenomic data alone are inadequate for predicting functional regulatory variants in a given cell type.

While there is a clear excess of *de novo* variation in coding sequences in ASD and other neurodevelopment disorders, substantial additional burden is estimated to be borne by other elements, such as transcriptional (8) and translational regulators (*e.g.*, splicing 5’/3’ untranslated regions; UTRs) (14,15). These regions function to regulate transcript stability and miRNA interactions (16); they may also mediate nuclear retention/export of transcripts in the brain, which has only recently been explored (17). Coding variation carries a minority of heritable risk for schizophrenia, bipolar disorder, and attention deficit hyperactivity disorder (18). The occurrence of most disease-linked variation in the least-well understood features of the genome/transcriptome thus obstructs understanding of disease biology.

Collectively, these two problems necessitate **high-throughput assays with functional readout** for putative regulatory regions and variants. Such assays enable identification of functional variants and of the biological and environmental contexts in which they act. This knowledge, in turn, can begin shaping hypotheses regarding the shared mechanisms by which seemingly disparate genetic factors converge on shared phenotypic endpoints.

Here, we will primarily discuss MPRAs in terms of their potential for high-throughput parcellation of genetic studies. This technology relies on pairing genomic features (*e.g.*, an allelic pair of a variant-containing region) to a reporter gene bearing unique, transcribed barcodes, allowing an RNA-level readout of features’ activities in a multiplex setting (19,20). This approach can be flexibly implemented allowing study of transcriptional regulators (*e.g.*, promoters), as well as splicing, protein translation, and post-transcriptional regulators. Critically, the potential for MPRAs to dissect roles of variants from neuropsychiatric genetic studies is far from fully realized. Here, we aim to illustrate that MPRAs are a methodologic “low-hanging fruit” for neuropsychiatric genetics. In the first part of this review, we discuss the importance of cellular, biological, and environmental contexts in the design and execution of MPRA, exemplify uses of the approach to date, and discuss complementary/follow-up methods to further validate functional variants nominated by MPRA. In the second part, we discuss multiple features of MPRA that are well-suited to parsing polygenic architecture—both for dissecting linked blocks of multiple functional variants, and for identifying convergent variant mechanisms across the genome.

#### Part 1: MPRAs for Identification of Sequence Variants with Functional Consequences

MPRAs **quantify activity of putative regulatory elements by coupling them to a reporter gene and counting transcribed, element-specific tags (“barcodes”), using sequencing to determine the ratio of (expressed RNA barcode)/(delivered DNA barcode).** This experimental framework has been applied to human splicing (21–23), RNA editing (24), protein translation (25), UTRs (26–30), and, most broadly, transcriptional (*i.e., cis-*) regulators. MPRAs thus offer a flexible framework to study regulatory phenomenon, including transcription factor (TF) and RNA binding protein (RBP) actions, transcript stability, and ribosome occupancy. MPRAs have been most broadly applied to explore and computationally model “regulatory grammar” of transcriptional regulators: how sequence features such as binding motifs, their abundance, and arrangement affect regulatory capacity (31– 38). More recently, these approaches have been applied to identify the transcriptional consequences of SNPs and rare variants (39–45).

As shown in **Figure 1A-B**, a canonical ‘enhancer’ MPRA utilizes a promoter with candidate elements either immediately upstream or in a 3’UTR (STARRseq) (same approach as **Figure 1D**) (46). Each sequence element is paired to multiple, unique barcodes, which are transcribed into UTRs and sequenced as quantitative readout. Expression—representing transcription or RNA stability—is typically measured as the counts of RNA barcode per encoding DNA barcode (**Figure 1G**). To define active or differentially active elements, expression levels can be normalized to *e.g.*, a minimal-promoter only set of barcodes (31,34,37,47–49), compared between alleles (39–45), or compared to shuffled parent sequence(s) (32,37)

**Figure 1.**
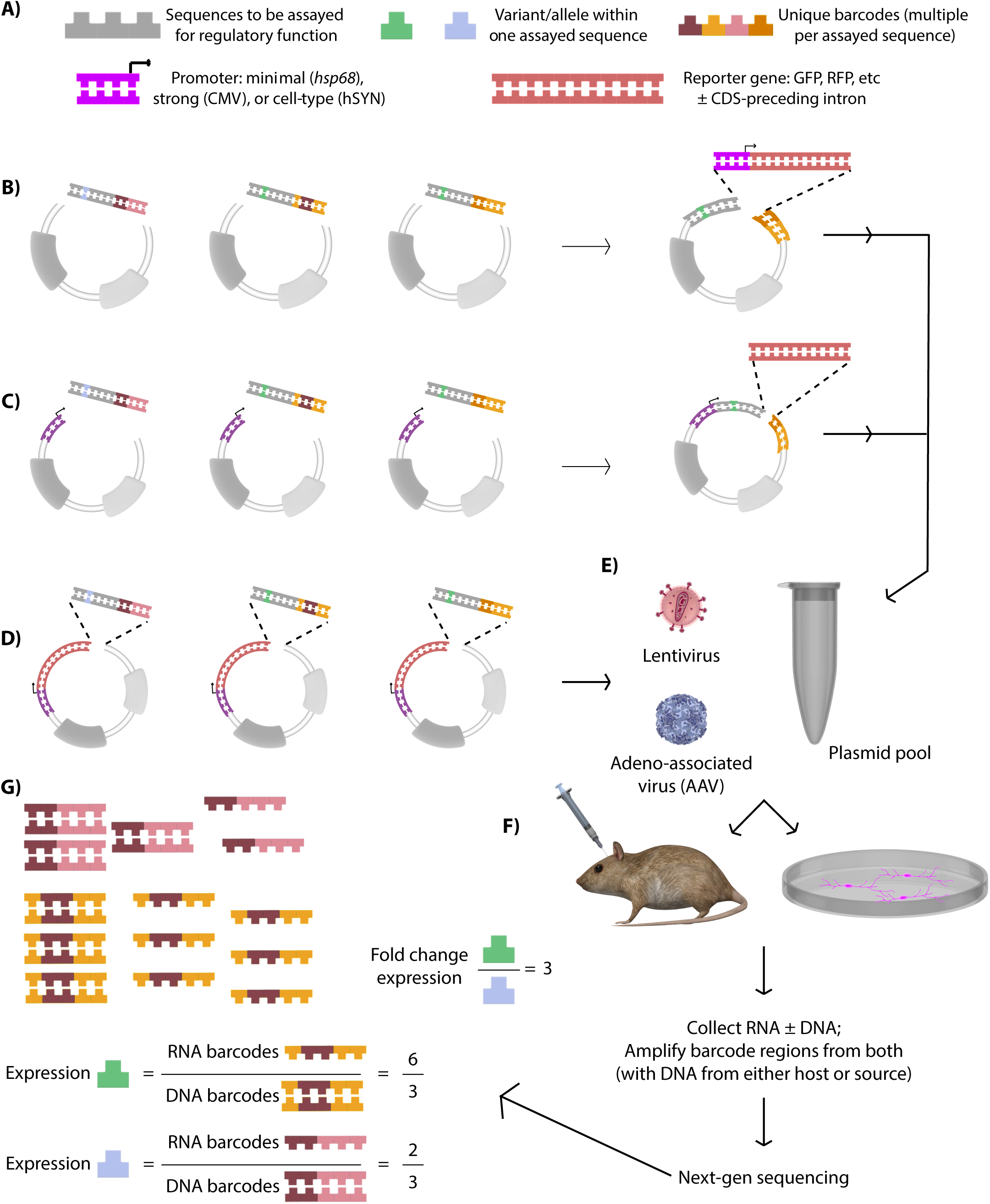
Cloning, Delivery, and Analysis Approaches in MPRA. **A)** Visual key for subsequent panels. **B)** Enhancer MPRAs most commonly clone a pool of custom oligonucleotide pools containing sequences to be assayed, each paired to multiple unique barcode sequences, into a vector. This plasmid pool is collected and a promoter-reporter cassette is subsequently inserted between the elements and barcodes, such that the element of interest is upstream of a promoter and the barcode is in a 3’UTR. In the case of STARR-seq (not shown), the paradigm in panel **D** is instead used, with the *cis-*regulatory element acting as its own barcode. **C)** 5’UTR assays likewise use a two-step cloning assay, placing elements immediately *down*stream of a promoter. A reporter gene itself is inserted between the element and barcode. **D)** 3’UTR assays place both the elements of interests, immediately adjacent to barcodes, downstream of a promoter-reporter in a single cloning step. **E)** Sequence pools from approaches B-D can then be packaged into AAV or lentivirus (if in a compatible vector), or used for assay as plasmid directly. **F)** The plasmid or viral MPRA library can be delivered to cells in culture or in vivo tissue. After transfection/transduction, nucleic material is collected to generate sequencing libraries to quantify expression of the delivered elements. **G)** MPRA analysis centers on taking the ratio of RNA/DNA counts (or counts per million), represented by the sequence fragments at top left, as a measure of expression.

MPRAs also enable study of post-transcriptional regulatory elements. As shown in **Figure 1C and 1D**, the same architecture and RNA/DNA expression metric can be used to assess UTR effects on transcript stability. UTR MPRAs have yet to be implemented to study regulatory variants directly, but have been used to study the regulatory grammar of the ASD/ID-implicated CELF proteins and related RBPs (26,50,51), features conferring transcript stability via 3’UTRs (27,28,30), and 5’UTR influences on translation (52) and prediction of functional 5’UTR variants (29). Across enhancer and UTR MPRAs, several key forms of disease-associated noncoding variation can be assessed for functional consequences using a variety of model systems and delivery approaches (**Figure 1E-F**).

### MPRAs Enable Identification of Functional Regulators and Variants In Specific Cellular Contexts

Perhaps the most exciting—if underappreciated—property of MPRAs is the ability to assay regulators of interest in the relevant cell types and conditions. Functional elements both define—and are defined by—a given cell type’s unique milieu of expressed TFs, chromatin modifiers, miRNAs and RBPs, which in turn specify regulatory element functionality. The importance of cell type is strikingly illustrated by a *cis*-regulatory MPRA of random sequences performed in parallel using U87 glioblastoma and neural progenitor cells. Comparison of the most active enhancers across the two cell types yielded entirely different motifs and sequence features correlated with expression (49). This **sensitivity to cellular context** highlights that **MPRAs can be designed to assay and compare genetic regulation in cell types known—or predicted to—mediate disease.** This also highlights the fact that careful consideration needs to be given to the appropriate cellular context when designing assays for psychiatric genetics: the array of variant-interacting TFs and RBPs expressed in neural cells may not be present in convenient cell lines (*e.g.*, HeLa or HEK-293).

While cell type overwhelmingly influences outcomes in regulatory assays, additional conditions could equally alter outcomes (**Figure 2**). Age, sex, pharmacology, and environment (*e.g.*, stress)—all, like cell type, can shape or reshape regulatory activity. This capacity of MPRAs has recently been applied in neurogenetics—specifically in determining the temporal patterns of regulatory element activity over the course of differentiation of human neural progenitor cell (NPC) differentiation to neurons (53).

**Figure 2.**
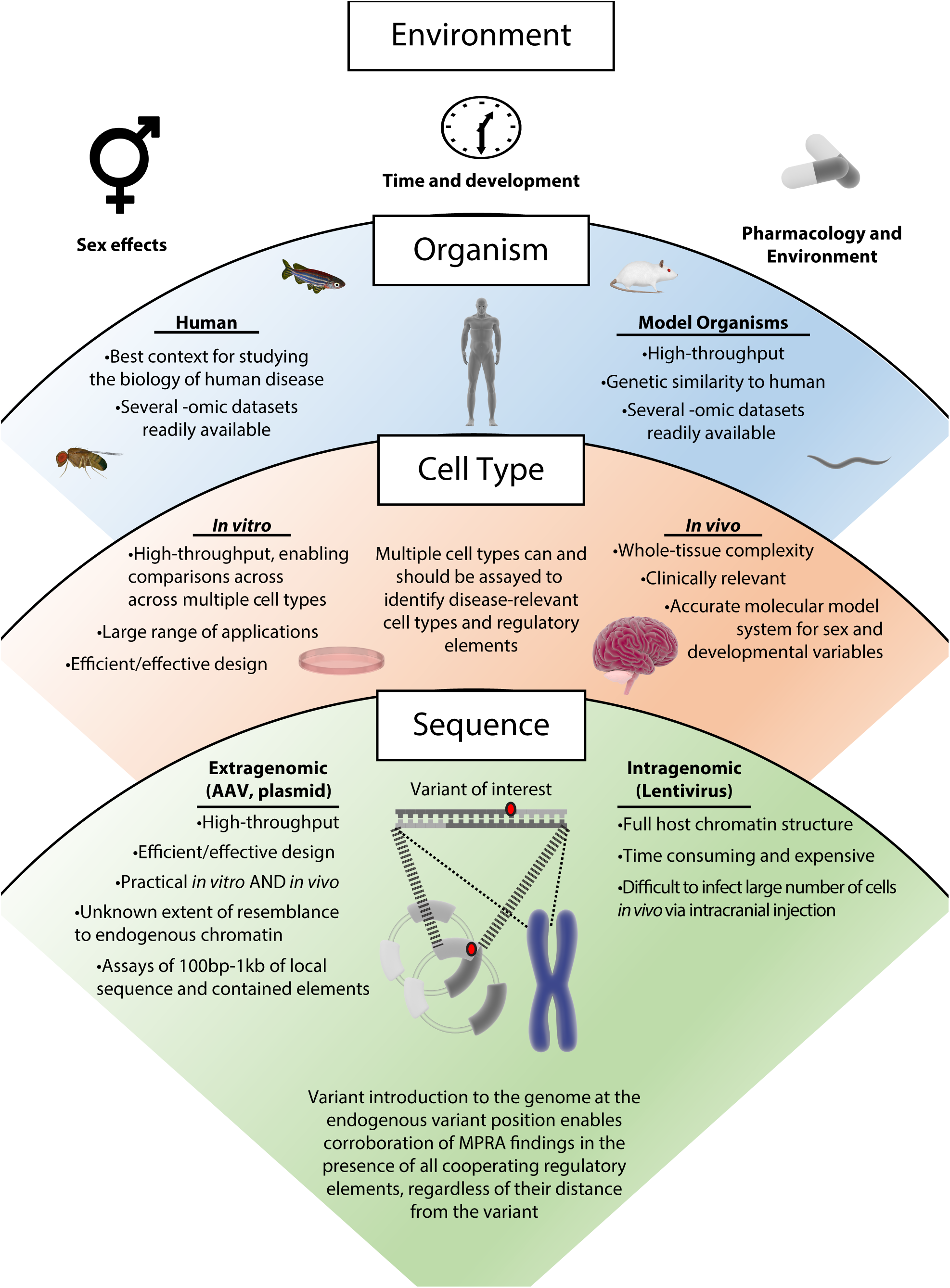
Regulatory assays are influenced by a range of conditions, from environment to sequence context. The range of conditions that influence regulatory assays (from top to bottom) starts when considering the environment, *e.g.*, sex, time, and pharmacology. These parameters have the potential to affect various –omic profiles in a given system. The next level of consideration is the organism, which can include human-derived tissue or one of the many model organisms. Human genomic context is ideal for studying the biology of human disease – though a comparatively limited scope of techniques for human-derived tissues exists. Next, one should consider the selected cell type(s) and whether to assay *in vitro* or *in vivo*. Each of these provides a unique set of benefits, and one approach can be used to verify findings from the other as a form of validation (45,74). In the case of modeling the brain and psychiatric genetic variants, cell type-specific/enriched MPRAs *in vivo* constitute the highest-fidelity model of variant effects by accounting for regulatory effects of endogenously interacting cell types. Lastly, the sequence context will be influenced by the delivery method, which results in transcription from extragenomic or intragenomic MPRA DNAs. In either case, only limited length of sequence surrounding a feature of interest is preserved (*e.g.*, in ∼120bp tiles of genomic sequence in custom oligonucleotide cloning, or ≤ 1kb in clone-and-capture methods), preventing assessment of any interactive effects from elements further away. (A recent study suggests that size of a tile negatively correlates with reproducibility of expression driven compared to that driven by ∼120bp tiles, emphasizing the importance of this consideration (93)). While AAV-transduced episomes gain histones (105) and chromosome-like nucleosome spacing (106), it is unknown whether gene-regulatory histone marks on these episomes mirror those of endogenous regulatory chromatin. For these reasons of both local sequence context and chromatin context, we suggest corroboration of MPRA findings in native genomic settings, by, for example, introducing the variant to the genome of a cell line using CRISPR methods.

In addition to context-specific element activity, MPRAs can also identify context-specific functional *variants*. This follows from findings that the influence of genomic variants on epigenomic marks is also cell type dependent, even within a developmental lineage: approximately 80% of SNPs with allelic associations to chromatin accessibility in human NPCs or neurons only carry such an association in one of these two cell types (54). Thus, experimental study of putative disease-associated variants requires firm hypotheses on where (cell type), when (development/differentiation), and how the regulators are expressed/active and biologically relevant. For example, a hypothetical study of regulatory variants affecting action of a candidate ASD gene like *CHD8*—expressed most highly in early development (55)—using mature neurons may not detect any effects because, while matched on cellular context, the temporal context is inappropriate for examining the candidate gene. The importance of cell type in identifying regulatory variants is explicitly highlighted by the enrichment of GWAS SNPs with tissue-specific associations to gene expression (56,57). **MPRAs are thus poised to be applied to study of cell type-specificity of regulatory variation both *in vitro* and, as we will demonstrate, *in vivo*.**

Work toward assessing and predicting the effects of biological context on genomic variation has been recently illustrated in computational genomics. The Critical Assessment of Genome Interpretation 5 (CAGI5) consortium performed an MPRA of saturation-mutagenized human regulatory elements and disease-associated promoters in numerous cell types. A subset of these data were provided to analysts, who were then challenged to computationally predict functionality and effect sizes of held-out variants (58). The end project focused on identifying the most informative datasets (*e.g.*, chromatin modification data from ENCODE) for cell-type specific regulatory variant prediction. A key finding was that the most informative datasets often came from the same or similar cell types as that for which predictions were made.

Complimentarily, MPRAs focused on neuropsychiatric disorder associated variation stand to benefit *from* high-information datasets by aiding variant prioritization for assay inclusion. Several datasets on synthetic UTRs (27,29), RNA binding proteins (59,60), and postmortem human brain multi-omics (61– 70) have become available in recent years. Integrative computational analyses have brought these datasets together predict functional variation in SCZ, bipolar disorder, and ASDs (71,72); however, these predictions have not yet been systematically tested. These constitute high-priority candidates for experimental validation by MPRA. Finally, inclusion of MPRA data with regulatory-epigenomic annotations improves machine learning predictions of functional variants (73). As such, MPRAs and omics form a symbiotic loop whereby MPRA results refine regulatory omics data, improving variant prioritization for future computational and functional investigations.

In spite of their potential, few MPRAs have examined regulatory variation while considering both cell type and –omic predictions. Tewhey, et. al (41) implemented an MPRA in human lymphoblastoid cell lines (LCLs) to identify functional regulatory variants among nearly 30,000 eQTL SNPs from LCLs, thus maintaining an appropriate cellular context for testing the discovered variants. Over 3,400 active regulatory sequences were identified, including 850 activity-modulating variants (24%), detectable even though effect sizes generally were under 2-fold. More recently, Choi, et. al (74) prioritized over 800 SNPs—guided by fine mapping and epigenomic signals—from 16 melanoma GWAS loci, then assayed the variants for transcriptional-regulatory activity in melanocytes *in vitro*. Candidate variants with concordant eQTL signal in independent melanocyte data were prioritized for follow-up, ultimately enabling their experimental demonstration of biophysical (TF binding), molecular (target gene expression), cellular (growth rate), and *in vivo* (melanoma rate in target over-expressing zebrafish) variant mechanisms. These experiments exquisitely illustrate MPRA’s capacity for sensitivity, context specificity, and high discovery rates, especially when integrating both association data and multi-omic annotations to nominate variants.

However, the ability of MPRAs to address clinically-relevant questions has, to date, largely been restricted by *in vitro* implementations, which can neither model the complex cell-type interactions of tissues—especially those of the brain—nor can they model tissues’ full breadth of sex/developmental, environmental, or pharmacological complexity **(Figure 2)**. Critically, even the most complex *in vitro* models of brain tissue—organoids—lack almost 50% of cell type-specific accessible chromatin regions found in their *in vivo* counterpart (fetal brain) (75), suggesting that *in vitro* MPRAs alone cannot dissect functional variation in neuropsychiatry. As preclinical research tools, *in vivo* MPRAs would greatly expand the ability to assess regulatory activity in native cellular and physiologic contexts. Two *in vivo* MPRA studies to date from Shen, et. al, confirm that MPRA is plausible for the brain (mouse cortex) *in vivo.* In the first, a library 45,000 barcodes corresponding to human brain open chromatin regions were packaged in adeno-associated virus 9 (AAV9) and transduced into adult mouse brain by stereotaxy (45). Consistent with open chromatin as a proxy for transcriptional regulation, many of the strongest accessibility peaks also drove gene expression in the mouse brain. In the second study, a single candidate SNP from a locus associated with bipolar disorder was prioritized by epigenomic data and ultimately demonstrated to have functional consequences; the two alleles of this sequence region were paired to 20 barcodes each, delivered by electroporation of embryonic mouse brains, which were explanted, maintained *ex vivo* for two days, and then assayed (44). While the two studies represent a major step toward assessing gene regulation in intact tissues, the barcode readout nonetheless comes from ‘bulk’ tissue RNA, leaving a limitation (albeit acknowledged) regarding cell-type effects on regulation.

### The frontiers of context-driven MPRA: A Proof of Principle *In Vivo*, Cell-Type Specific MPRA

To determine whether MPRA can be used to assay cell-type specific gene regulation in the *in vivo* setting of the brain, we generated four barcoded sets of AAV-compatible plasmids, containing a human promoter from pan-cellular (*PGK2)* (76), excitatory-neuronal (*CAMK2A)*, or astrocytic (*GFAP*) (77) genes, or a minimal promoter (mouse *Hsp68*) (34) alone.

First, *CAMK2A, GFAP*, and minimal-promoter-only (*Hsp68)* viruses were individually injected into P1 mice to give widespread transduction of the brain (78). Brains were collected and sliced at P21 to examine cell-type expression patterns of dsRed under each promoter, confirming the neuronal, astrocytic, and non-specificity of these three promoters, respectively (**Figure 3**).

**Figure 3.**
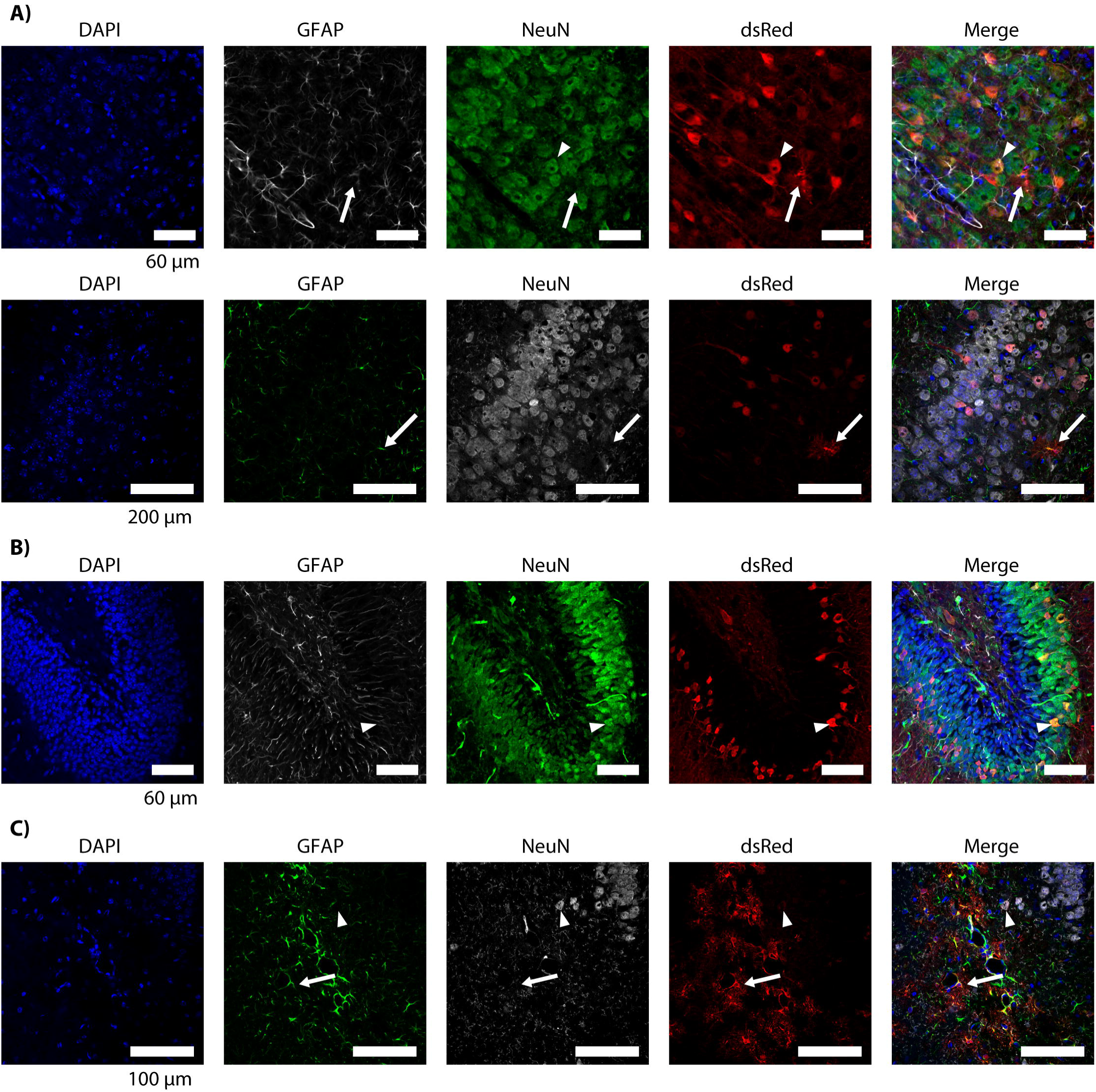
Immunofluorescent confirmation of cell type-specificity of promoters delivered by AAV9. All images are from mouse brain transduced with AAV9 at P2 and collected at P27. The AAV delivered contained dsRed under the control of the hsp68 minimal promoter, with or without an additional promoter upstream. Astrocytes were labeled with anti-GFAP and neurons with anti-NeuN antibodies. White arrowheads indicate examples of transduced neurons; white arrows indicate example transduced astrocytes. **A)** *Hsp68* alone drives dsRed expression predominantly in neurons in both the cortex (top row) and hippocampal granule layer (bottom row). Only three astrocytes (two shown) were found expressing dsRed across three examined slices of transduced forebrain. **B)** The *CAMK2A* promoter exclusively drove neuronal expression, shown in the hippocampal dentate gyrus granule layer. **C)** The GFABC1D promoter (subregion of the *GFAP* promoter) drove predominantly astrocytic dsRed expression, shown near the granule cell layer of the hippocampus. Expression is predominantly in astrocytes, such as those ensheathing blood vessels as shown, with rare, comparatively low expression in neurons.

Subsequently, the viruses were mixed at a titer of 1:1:1:1 and injected into the P1 brain of SNAP25-TRAP mice (n=2), from which neuronal RNAs can be selectively enriched using the TRAP method (79,80). At P28, brains were collected for TRAP to obtain neuronal and brain-wide RNAs for barcode sequencing alongside DNA from the viral mix delivered.

As expected, the neuronal TRAP fractions relative to the whole brain homogenate displayed increased RNA/DNA ratio (*i.e.*, expression) of the *CAMK2A*-driven barcodes, and relative depletion of the astrocytic *GFAP*-driven barcodes (**Figure 4F**), indicating that we were able to successfully identify cell-type differential activity of these regulators. The calculated expression on a per-barcode basis was highly replicable between the two animals (R^2^ > 0.99) (**Figure 4D and 4E**). In all, this experiment demonstrates that MPRAs are not restricted to tissue culture models and have the capacity to be executed in specific cell types in model organisms, enabling functional assessment of gene regulation in higher-fidelity developmental, hormonal, and physiologic contexts.

**Figure 4.**
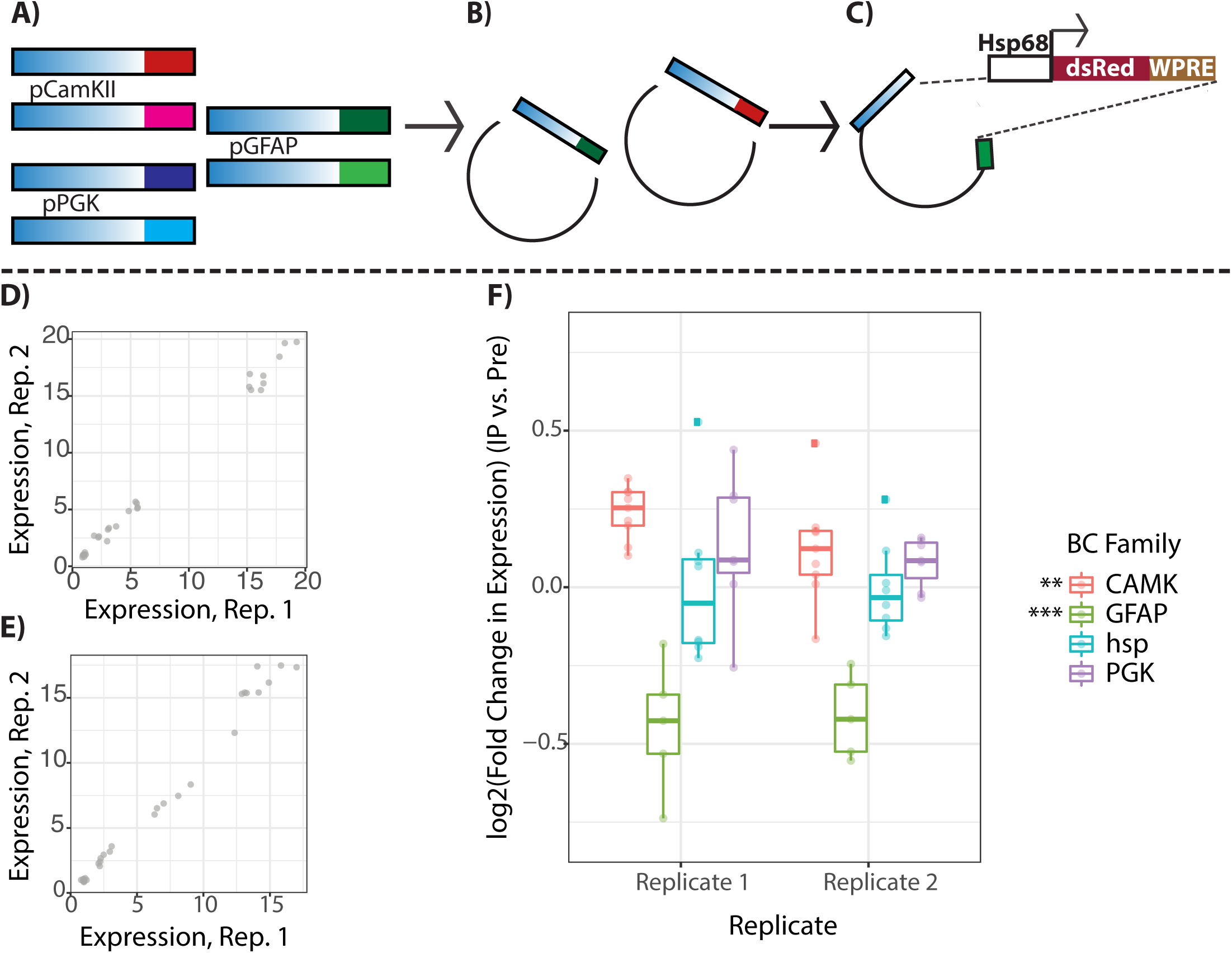
Design, replicability, and cell type-specificity of a small-scale, proof of principle *in vivo* MPRA. **A-C)** Design of the MPRA plasmids. For each promoter, multiple barcodes were PCR coupled to each promoter (**A**), inserted into plasmid (**B**), and an hsp68-dsRed construct subsequently inserted (**C**). Plasmid pools were created subsequently packaged in AAV9 for each promoter separately. The final library consisted of 32 barcodes across the four promoters. **D-E)** Mouse brains transduced with an equal-titer mixture of all 4 promoter AAV9s and subjected to neuronal (SNAP25) TRAP showed strong replicability of barcode-wise expression in both the neuronal-enriched TRAP RNA fraction (**D**, R^2^=0.9975) and total tissue (**E**, R^2^=0.9927). **F)** Expression across barcodes in each replicate confirmed that the TRAP fraction was enriched for *Camk2a*-coupled barcodes (neuron-specific) and depleted for GFABC1D-coupled barcodes (astrocytic) as expected. ** *p* ≤ 5 × 10^−4^, *** p ≤ 5 × 10^−8^

### Complementary methods in high-throughput study of DNA and RNA regulatory elements

A primary limitation of MPRA is the inability to test regulators in their endogenous genomic position and sequence context. This prevents identification of target genes (for transcriptional regulators or their variants) and confirmation of a variant or element’s relevance in its native genomic context. Sequence-specific targeting methods using CRISPR/Cas9 have enabled several additional techniques for probing molecular and cellular effects of regulatory variation, with the caveat that, in contrast to MPRA, these techniques do not currently allow for the multiplexed study of post-transcriptional/translational regulators. Nonetheless, these techniques complement MPRA well in that they enable study of the host genome directly. *Perturb-seq* (81) combines genewise perturbation by CRISPR with single-cell RNA-seq to identify gene sets dysregulated by loss of function of each candidate gene. These have, for example, been used to discover co-transcribed gene networks involved in neuronal remodeling (82), and for *in vivo* assessment of a set of genes discovered to harbor *de novo* loss of function mutations in ASDs (83). CRISPR editing has been used *in vitro* to assess single transcript variant effects by comparing reference and allelic RNA and genomic DNA abundances in edited cultures(84). While this approach currently requires a separate culture of cells for each assayed variant, it still enables practicable validation/follow-up of MPRA candidates by the dozens using, *e.g.*, 96-well plates. To our knowledge, such assays have not been conducted at a genome-wide scale in psychiatric disease, but have been used to identify genes that alter expression of the Parkinson’s-associated *PARKIN* (85).

*Cis-*regulatory MPRAs do not disentangle target gene(s) from functional regulators. Fortunately, CRISPR-derived methods using a mutagenically-’dead’ Cas9 (dCas9) conjugated to a transcriptional activator or repressor allow an experimenter to target and potentiate or repress endogenous genomic regulatory elements (CRISPRa and CRISPRi, respectively) to assess altered gene expression and other outcomes. These technologies are already online in state-of-the-art human neuroscience models: a recent CRISPRi study knocked down over 2000 genes by targeting their promoters in iPSC-derived excitatory neurons, defining gene roles in their survival, differentiation, and proliferation. Like MPRAs, such assays retain the ability to assess cellular context—in this case, a subset of cells were co-cultured with astrocytes, revealing that knockdown of certain genes had different consequences on neuronal survival in co-culture conditions (86). The cell-type specificity of CRISPRi has also been leveraged to study ASD-associated gene knockdown effects in an etiologically relevant cell type, NPCs (87). A recently introduced extension of CRISPRi targets intergenic regulators along with fluorescent *in situ* hybridization against genes in the same chromodomain to identify regulator-target pairs. Fluorescence-intensity sorting into bins and subsequent RNA sequencing of stained cells then could be used to pair regulators (via guide RNA sequence) and target genes (altered fluorescent intensity associated to a guide RNA’s presence) (88). While this assay was performed in K562 cells, it is not hard to envision its extension to neural cell types *in vitro* or *in vivo*. Altogether, CRISPR-based follow-up of MPRA candidates to define target genes and verify of genomic activity of regulators/variants will be key to developing insights in psychiatric genomics.

#### Part 2: MPRAs as an avenue to dissect multiallelic and polygenic mechanisms of neuropsychiatric traits

While MPRAs cannot intrinsically scale up to functional demonstration of cell-, tissue-, or behavior-level phenotypes, they have the potential to provide key information to guide molecular hypotheses for how these higher-order phenotypes emerge from large sets of regulators and/or their target genes. We will divide this discussion into a brief examination of *cis* variation—that is, the study of multiple elements in the same linkage block —and examination of variants in *trans* space—that is, defining shared and recurrent features among MPRA-nominated functional variants across the genome that may collectively underlie large portions of polygenic disease risk.

### The utility of MPRAs for parsing linked variation

One challenge in parsing loci implicated in common variant association studies is that multiple variants in the region are equally statistically associated, and thus equally plausible functional mediators of risk. Indeed, in statistical genetics, repeating a GWAS or eQTL association analysis after conditioning on the lead SNP within a block often reveals one or more additional independent variants associated with traits or gene expression, respectively (*e.g.*, (70,89,90)); in fact, nearly half (∼8,000) of brain expressed genes have ≥1 conditional eQTL (91)Reporter assays have been used to systematically evaluate such linked sets of variants. One disease-oriented, small scale example used epigenomic data to identify 16 enhancers spread over a few hundred kilobases near the *Ret* gene, known to be downregulated in Hirschsprung’s disease (failure of terminal colonic nerves to form *in utero*). The 16 putative regulators were assayed for allele-differential activity in neuroblastomas to model regulatory activity in the disease-relevant cell type, neural crest cells. These were likewise assayed with transient expression in mouse embryos driving a LacZ reporter to identify the crest-relevant regulatory sequences. They also validated roles for regulator-binding TFs via siRNA knockdown. In all, this identified three functional SNPs in linkage disequilibrium(92). While this was a small scale example, it is easy to imagine how MPRAs might be used to dissect the large and numerous blocks of common variation associated with psychiatric disease (3,4).

While the *RET* SNPs were several hundred kilobases apart, another challenge is with variants in very close proximity, potentially within the same regulatory element. While oligo synthesis is limited in length, MPRA-based assay of such variants spanning up to 700bp is now possible with use of PCR amplification of oligos with uniquely complimentary ends (93). Likewise, ‘capture-and-clone’ MPRA designs, often using STARR-seq architecture to simplify cloning, fragment genomic DNA or capture e.g. ChIP-seq DNA fragments to test larger fragments than attainable via oligonucleotide synthesis(45,94–96). Altogether, MPRAs allow for discovery of multiple functional variants per linkage region, as well as close-proximity discovery and disentanglement of multi-variant regulatory effects.

It is worth mentioning that MPRA could be of utility in the longer term as a technical tool to improve GWAS methods. Unusual linkage patterns in genomic regions such as the major histocompatibility complex confound localization of trait associations, leaving the region excluded from analyses or nebulously associated in others—over 200 traits (64), with MDD among them (65). Comprehensive MPRA of variants from this region across a broad panel of cell types would enable identification of cell/tissue-specific functional variants, which could then be placed on SNP genotyping arrays, partially sidestepping linkage limitations in understanding this region by collecting genotype information *at* the functional variant positions. Such efforts at a consortium scale could likewise characterize functionality and properties of broader sets of both untranscribed and untranslated variants in the same manner in each of several cell types.

### The utility of MPRAs in identifying commonalities from variants across the genome

The most vexing question that remains after individual functional variant mechanisms are elucidated is how variants *collectively* contribute to phenotypic risk. MPRAs provide several ways to begin addressing this question: 1) by identifying shared regulatory features across several functional risk variants; 2) identifying functional modules enriched for genes dysregulated by disease-associated variants; 3) by providing functional annotations to variants that can be integrated into computational genomic approaches; and finally, 4) by enabling study of variant-by-environment interactions using the same MPRA libraries across conditions.

Firstly, MPRA experiments running the gamut from basic regulatory genomics to human traits and variation have defined scoping ‘regulatory grammars’ of assayed contexts. Identification of functional variants in the MPRA setting enables similar establishment of the “regulatory grammar” of a trait or disease. Functional variants identified by MPRA across several UTRs may feature a specific RBP’s binding site, for example. Likewise, variants associated with a trait could be more likely to fall in particular TF binding sites or be enriched cell-type specific marks of genomic regulation. Evidence of this convergence is seen in *de novo* variants associated with ASD: several distinct variants disrupt binding sites for a single TF, *NFIX* (97). Such an approach was taken in a recent MPRA of lupus-associated SNPs, where identified functional variants were intersected with TF ChIP-seq datasets from pertinent cell types (leukocytes), identifying sets of recurrently disrupted TF binding sites (98). Assays of downstream consequences of variation also confirm biological convergence across association loci. A four-gene-target CRISPRi/a assay revealed that schizophrenia risk genes act synergistically via shared influence on synaptic activity, and concurrent alteration of expression of all four genes results in a cellular transcriptome more accurately reflective of postmortem schizophrenia brain tissue(99). For both rare and common variants, identifying common regulators among risk genes provides information which can refine predictions of disease-related cell types based on their expression of such TFs, RBPs, or epigenomic marks.

Secondly, genes and gene networks affected by statistically associated variation are often predicted using MAGMA (100), which in essence scores genes based on proximity to an associated variant and its linkage partners. Resulting gene sets are subjected to analyses such as Gene Ontology enrichment or are examined for enrichment in WGCNA (co-transcriptional) networks from candidate tissue types to identify pathways and mechanisms on which these genes converge. While its use is ubiquitous in genomic studies, standard MAGMA gene association statistics for psychiatric disorders only modestly correlate to those from a tissue-specific, chromatin configuration-aware modification of MAGMA (101), suggesting that biological hypotheses from MAGMA gene sets may miss disease-associated genes in brain. Being able to refine implicated genes by functional validation using—or in follow-up to—MPRA will help to benchmark such approaches and refine prediction convergence with ‘truly’ dysregulated candidate genes.

Thirdly, epigenomic data alone is not comprehensively predictive of active regulators. However, well-informed analyses of human genetic findings rely heavily on such annotations to convert associations into biological hypotheses. Critically, these epigenomic data—unlike MPRA data—can be collected from postmortem human tissue. However, a symbiotic loop is possible wherein verified regulators implicated by MPRA could be used to train and improve annotation of epigenomic algorithms —which in turn could be assessed by MPRA and used as another training set. Refined interpretation of epigenomic data and variant-expression association can then improve myriad analyses, such as variant enrichment in classes of genomic features, as well as disease gene identification. Other high-order analyses, such as TWAS (102) and Predixcan (103) intersect gene expression QTLs (eQTL) with trait-associated variants to predict expression differences between cases and controls, thus identifying dysregulated gene sets. MPRA data can disentangle which eQTL SNPs are truly functional from those associated only due to LD. Layering in MPRA data as weights or qualifiers of SNPs included in these analyses ought to refine disease-gene set associations much as discussed above. Altogether, MPRA can serve to refine both epigenomic and genic definitions of truly causal disease features.

Finally, the context-specificity of MPRA (**Fig 2**) represents a newfound ability to assess variant effects on gene regulation *en masse* under different biological and environmental contexts, including with *in vivo* models. While issues of convergent disease effects across genes and regulators are indeed complex, environmental effects—perhaps most canonically, stress—on these regulators are questions at the forefront of understanding polygenic risk in neuropsychiatric disorders. Pharmacologic variables have been successfully tested in MPRA, namely in the identification of glucocorticoid-responsive (104) and p53-responsive (96) regulatory elements. MPRAs could further be layered with concurrent gene perturbations (*e.g.*, knockdown of a putative regulator) or cell culture conditions for *in vitro* identification of variant-environment interactions. However, the most exciting function of MPRAs may be the opportunity to study disease-associated factors that cannot be (well) modeled *in vitro*, such as sex or stress, by moving these assays *in vivo*. Such studies can clarify sets of variants which depend on specific exogenous (*e.g.*, stress) or endogenous (*e.g.*, sex) contexts. Such variants could indicate molecular mechanisms behind conditional disease risk in a disorder such as depression.

## Conclusion

MPRA presents unique opportunities to dissect polygenicity of psychiatric disorders via simultaneous identification of functional variants across identified risk space. Beyond the primary benefits of identifying ‘true positive’ functional variants in specific biological and environmental contexts, MPRAs stand to rapidly broaden, deepen, and refine hypotheses and mechanisms of both noncoding disease risk and of gene-regulatory architecture itself.

## Supporting information

Supplemental Table 1: Antibodies Used for Immunofluorescence

Supplemental Text: Experimental Methods (In Vivo TRAP-MPRA)

## CONFLICT OF INTEREST

No authors declare a conflict of interest.

## ACKNOWLEDGEMENTS

This work was funded by NIMH grants 1F30MH1116654 and 1R01MH116999, Simons Foundation 571009 and NHGRI grant 1R01HG008687. The *Camk2a* promoter was derived from a plasmid generously gifted by Karl Deisseroth to Washington University’s Viral Vector Core. We would like to thank Sergej Djuranovic, Ph.D., Barak Cohen, Ph.D., and Cohen lab alumni Dana King, Ph.D., and Brett Maricque, Ph.D. for their collaboration and guidance designing, adapting, and analyzing MPRAs applied to neuropsychiatry. We would also like to thank Idoya Lahortiga, Ph.D. and Luk Cox, Ph.D. curators of Somersault1824 (https://www.Somersault1824.com), for their open-access, Creative Commons BY-NC-SA 4.0 licensed libraries of high-quality biomedicine graphics (especially those from Graphite Life Explorer, ePMV, and Eyewire), adapted for figures in this review. Finally, we would like to thank Mike Vasek for his assistance in editing this manuscript.

## Notes

#### Summary of Updates

Added discussions of A) recent, highly-relevant literature supporting principles behind MPRA, and B) additional UTR-focused MPRAs. Corrected several major copyediting issues in the early part of the text, which obscured the meaning of information critical to understanding the principles of MPRA (and thus the rest of the paper).

