## Supplemental Text: Experimental Methods (In Vivo TRAP-MPRA) for "The Oft-Overlooked Massively Parallel Reporter Assay: Where, When, and Which Psychiatric Genetic Variants are Functional?"

**Supplemental text: *In vivo* MPRA methods**

*Animal Research Statement*

All procedures involving animals were approved by the Institutional Animal Care and Use Committee at Washington University in St. Louis, MO.

*Design and construction of minimal promoter-reporter-WPRE 3’ UTR cassette.*

A previously designed MPRA reporter consisting of a minimal *hsp68* promoter and dsRed-Express2 (1) was PCR amplified with a forward primer adding a 5’ *MreI* cut site and a reverse primer with 3’ overhang homologous to the 5’ end of the WPRE 3’UTR element. The WPRE 3’ UTR element was PCR amplified from a lentiviral plasmid encoding cyan fluorescent protein with the WPRE element (derived from plasmid FCIV) (2) with a forward primer containing a 5’ overhang homologous to the 3’ end of dsRed and a reverse primer adding a 3’ *PacI* cut site. These PCR products were cut and purified from a 1% agarose gel and subjected to 10 cycles of PCR without primer to allow overhangs to anneal and act as primers to create a contiguous sequence, followed by addition of hsp68-dsRed forward primer and WPRE reverse primer and 15 further PCR cycles to amplify the contiguous product. The PCR reaction was run out on a 1% agarose gel and the properly sized band (~1.5kb = ~1kb hsp-dsRed + ~500bp WPRE) was cut and purified. During Sanger sequencing (*described below*), proper assembly of the cassette was confirmed.

*Design and construction of cell-type-specific promoter MPRA library*.

Three promoters were PCR amplified with primers adding an *MluI* cut site to the 5’ end, and *MreI* and *PacI* cut sites to the 3’ end (for later insertion of the hsp-dsRed-WPRE cassette). Promoters were **1)** a 2.2kb human *Gfap* promoter region (3), **2)** a 1.3kb mouse promoter region of the excitatory, neuron-specific gene *Camk2a*, amplified from plasmid pAAV.CamKII(1.3).eYFP.WPRE.hGH (Addgene plasmid # 105622, a gift from Karl Deisseroth to Washington University’s Viral Vector Core); and **3)** a 521bp human promoter region of the constitutive, highly transcribed gene *Pgk2*, amplified from plasmid pRRLsinPGK-GFPppt (2). These PCR products were used as template for a second PCR, in which a reverse primer homologous to the added *MreI and PacI* cut sites was used. The reverse primer also contained an overhang consisting of a 9bp region of N’s, resulting in random sequences for use as barcodes, followed by a *SalI* cut site for insertion of the product into a plasmid backbone. For the minimal-promoter only condition, a single oligonucleotide consisting of all four cut sites, intervening bases to ensure cuttability, and a barcode (*MluI*-*MreI*-*PacI*-barcode-*SalI*) was ordered and PCR amplified. For all promoter PCR products, barcode identities were determined by Sanger sequencing after insertion of PCR product into vector (see below).

Plasmid JD386 (4), originally encoding *mTdTomato* under the *Gfap* promoter, was digested at *MluI* and *SalI* sites (products #R3198 and #R3138, New England Biolabs, Ipswich, MA, USA**)** to remove the promoter and *mTdTomato*, leaving an intact human growth hormone (hGH) poly-A signal sequence just downstream of the *SalI* site, as well as intact ampicillin resistance and inverted terminal repeats necessary for AAV packaging. The promoter PCR product was digested with the same enzymes. The plasmid digest was treated with antarctic phosphatase (AP*)* (NEB #M0289) and gel purified to isolate the desired backbone fragment. The digested PCR products and gel-purified, AP-treated backbone digest fragment were ligated using T4 ligase (NEB #M0202) at 16°C for 18 hours with vector:insert molar ratios ranging from 1:3 to 1:5. Ligations were then directly transformed into DH5α chemically competent cells (NEB #C2987H), outgrown for 45 minutes in a 250rpm shaker at 37°C, then plated on LB agar with ampicillin and allowed to incubate for 16-18 hours at 37°C. Individual colonies were used to incubate 1mL wells of LB-ampicillin liquid media in 96-well deep-well plates and shaken for a further 16 hours at 37°C. The liquid cultures were then used to generate two identical 96-well plates of 25% glycerol culture stocks, one of which was sent for Sanger sequencing. Sequencing in this round of cloning used primers upstream of the *MluI* site and downstream of the *SalI* site, allowing verification of the promoter sequence and identification of barcodes. Up to 10 clones per promoter were selected on the basis of having both expected sequencing results and unique barcodes. Equal volumes of each clone’s glycerol stock was then used to inoculate a 10mL LB-ampicillin liquid culture of each promoter’s “library 1” and grown at 37°C for 18 hours.

The “library 1” cultures were then mini-prepped (NucleoSpin Plasmid kit #740588.250 Macherey-Nagel, Düren, Germany) to isolate plasmids for insertion of the previously mentioned reporter cassette. Each library 1 plasmid pool, as well as the reporter cassette, was digested with *MreI* and *PacI* (Thermo Fisher #ER2021, Waltham, MA, USA; NEB #R0547) . As before, the plasmid digest was treated with AP and gel purified. The digested, AP-treated, purified plasmid fragment and digested reporter cassette were combined at a vector:insert ratio of 1:5 and ligated using T4 ligase as above. Ligations were directly used for transformation, plating, and single-clone 96-well Sanger sequencing and glycerol stocking as described above. Sanger sequencing was performed with the same primers as for Library 1; this confirmed the presence of the inserted promoter and for shorter promoters (minimal-onlyhsp68 minimal promoter and 5’ portion of dsRed (sequencing downstream from *MluI* site), as well as retention of the barcode and the presence of WPRE and the 3’ end of dsRed (sequencing upstream from the *SalI* site). Clones were selected for generation of “library 2” on the basis of present and intact promoter, minimal promoter, dsRed, and WPRE, and were only selected if the barcode identified was one of the barcodes from among the library 1 clones. This resulted in 5 to 9 eligible clones for each promoter condition. The clones’ glycerol stock wells were again used as inoculum for a 10mL LB-ampicillin starter culture, used to inoculate 500mL LB-ampicillin cultures for maxiprep (Plasmid Maxi kit (#12163), Qiagen, Hilden, Germany). The four maxiprepped “library 2” plasmid pools (one pool per promoter) were submitted to the Washington University Viral Vectors Core for packaging into AAV9.

*Neonatal mouse transduction with cell-type promoter MPRA library(ies); subsequent TRAP and immunofluorescence (IF)*

One litter of SNAP25-RPL10a-eGFP TRAP mice (described in (5)), back-crossed for >10 generations to wild-type C57BL/6j mice, was genotyped at postnatal day 2 (P2). GFP-positive pups received intracranial injection of a mixture of all four viruses mixed at an equal titer; an aliquot of the mixture was set aside for later sequencing for DNA counts during MPRA analysis. GFP-negative pups (3) were injected with either the *hsp68*-only virus, *Camk2a* promoter virus, or *Gfap* promoter virus to confirm cell-type specificity by immunofluorescence. All animals were injected with 2µL bilaterally of virus or viral mix into three coordinate pairs: four anterior to bregma (two anteromedial, and two ~1.5mm posterolateral to the former), and two medially midway between bregma and lambda.

At P17, the three GFP-negative pups were perfused with 4% paraformaldehyde in PBS, brains dissected and dehydrated, and sliced in the coronal plane into 40µm floating sections stored in PBS with 0.01% sodium azide for immunofluorescence. Antibodies are described in Supplemental Table S1. After incubation in secondary antibody, tissue was thrice washed in 5mL wells of PBS for 5 minutes. DAPI (Sigma-Aldrich D9542, St. Louis, MO USA) was present in the PBS used for the second wash to stain nuclei. Sections were then slide-mounted and imaged on a Zeiss LSM 700 (Zeiss, Germany) using a 40x oil objective.

At P27, the three surviving GFP-positive mice were sacrificed for individual biological replicates of TRAP, performed as previously described (6,7). Briefly, whole brain was separately homogenized from each mouse. 2.5% of the supernatant from the 20,000 *x g* spin of homogenate was collected as a “pre-IP” sample of RNA, i.e. RNA from all cell types in the tissue. The remaining supernatant was used for immunoprecipitation targeted at GFP to enrich for ribosome-bound RNA in SNAP25-positive cells (i.e., neurons) / to deplete RNAs from other cell types. Both pre-IP and IP RNA were QC’ed using an Agilent Tapestation’s high-sensitivity RNA assay (Agilent, Santa Clara, CA). Pre-IP RNA integrity number estimates (RINe) ranged from 7.8-8.2; IP RINe values ranged from 6.8-7.4. The IP from one sample had a RINe of 2.7; this sample (both pre-IP and IP) was excluded from the experiment for a final n of 2 biological replicates.

*RNA-seq library prep*

RNA samples were treated with the Turbo DNA-Free kit (Ambion #AM1907, Austin, TX, USA) to remove extant DNA. Sequencing libraries were prepared from RNA by performing a variation on the methods of *e.g.* (1,8,9) by using a reporter-specific primer, targeting the polyA signal sequence just 3’ to the barcode sequence (10), during first-strand reverse transcription with Superscript III Reverse Transcriptase (Invitrogen 18080044, Carlsbad, CA, USA). Double-stranded cDNA was then synthesized and amplified by PCR using Phusion HF (NEB #M0531) using the same reverse primer and a forward primer in the WPRE of the 3’UTR. These primers added unique cut sites allowing subsequent Illumina adapter ligation. To prevent sequencer errors due to homogenous sequence at the start of read 1 (3’ end), these adapters were of four different lengths to stagger the first base of the 3’ end read. Digestion, clean-up, ligation, clean-up, and final PCR with primers with partial homology to the adapter ends were used to add the remaining Illumina sequences and sample indices per (1,8,9,11). Samples were sequenced on a MiSeq System (Illumina, San Diego, CA, USA) at ~400k reads per sample, with ~900k reads for the DNA for accurate normalization of RNA/DNA levels.

*Sequencing analysis*

Barcodes were counted from read 1 sequences, allowing up to 3 mismatches in the 20bp upstream of the barcode and 0 mismatches within the barcode sequence itself. The number of reads mapping to each barcode were totaled and normalized to counts per million (cpm) with normalization for sequencing library size (in number of reads mapped) using EdgeR (12,13). Expression for a given barcode was then calculated as the ratio of (cpm RNA / cpm viral DNA) for each RNA sample (IP and pre-IP RNA from each brain). Expression values were normalized to the within-sample mean expression of the minimal promoter (*hsp68*) alone by taking expression (BC in sample) / expression (mean(*hsp68* BCs)) ). By normalizing within sample (i.e., pre-IP or IP), expression is thus normalized to activity among the cell types and proportions comprising the RNA sample. Significance was calculated by performing repeated-measures ANOVA / linear mixed modeling. Each barcode group’s expression was implemented as a repeated measure, and modeled as the a dependent variable of RNA sample type and of a random variable for source tissue, thus: *expression ~ RNA.fraction + (1|mouse)* (11). For plotting and interpretation of barcode enrichment/depletion between biologically paired pre-IP and IP RNA samples, log2 fold-change in expression was calculated by subtracting log2(normalized expression in Pre) from log2(normalized expression in IP) for each barcode.

1. Kwasnieski JC, Fiore C, Chaudhari HG, Cohen BA (2014): High-throughput functional testing of ENCODE segmentation predictions. *Genome Res* 24: 1595–1602.

2. Li M, Husic N, Lin Y, Christensen H, Malik I, McIver S, *et al.* (2010): Optimal promoter usage for lentiviral vector-mediated transduction of cultured central nervous system cells. *J Neurosci Meth* 189: 56–64.

3. Lee Y, Messing A, Su M, Brenner M (2008): GFAP promoter elements required for region-specific and astrocyte-specific expression. *Glia* 56: 481–93.

4. Sakers K, Lake AM, Khazanchi R, Ouwenga R, Vasek MJ, Dani A, Dougherty JD (n.d.): Astrocytes locally translate transcripts in their peripheral processes. *Proceedings of the National Academy of Sciences of the United States of America* 114: E3830–E3838.

5. Dougherty JD, Fomchenko EI, Akuffo AA, Schmidt E, Helmy KY, Bazzoli E, *et al.* (2012): Candidate pathways for promoting differentiation or quiescence of oligodendrocyte progenitor-like cells in glioma. *Cancer Res* 72: 4856–68.

6. Doyle JP, Dougherty JD, Heiman M, Schmidt EF, Stevens TR, Ma G, *et al.* (n.d.): Application of a translational profiling approach for the comparative analysis of CNS cell types. *Cell* 135: 749–762.

7. Mulvey B, Bhatti DL, Gyawali S, Lake AM, Kriaucionis S, Ford CP, *et al.* (2018): Molecular and Functional Sex Differences of Noradrenergic Neurons in the Mouse Locus Coeruleus. *Cell Reports* 23: 2225–2235.

8. Kwasnieski JC, Mogno I, Myers CA, Corbo JC, Cohen BA (2012): Complex effects of nucleotide variants in a mammalian cis-regulatory element. *P Natl Acad Sci Usa* 109: 19498–503.

9. White MA, Myers CA, Corbo JC, Cohen BA (n.d.): Massively parallel in vivo enhancer assay reveals that highly local features determine the cis-regulatory function of ChIP-seq peaks. *Proceedings of the National Academy of Sciences of the United States of America* 110: 11952–11957.

10. Rabani M, Pieper L, Chew G-L, Schier AF (n.d.): A Massively Parallel Reporter Assay of 3′ UTR Sequences Identifies In Vivo Rules for mRNA Degradation. *Molecular cell* 68: 1083-1094.e5.

11. Rieger MA, King DM, Cohen BA, Dougherty JD (2018): CLIP-Seq and massively parallel functional analysis of the CELF6 RNA binding protein reveals a role in destabilizing synaptic gene mRNAs through interaction with 3’UTR elements in vivo. *Biorxiv* 401604.

12. Robinson, McCarthy D, Smyth G (2009): edgeR: a Bioconductor package for differential expression analysis of digital gene expression data. *Bioinformatics* 26: 139–140.

13. McCarthy DJ, Chen Y, Smyth GK (2012): Differential expression analysis of multifactor RNA-Seq experiments with respect to biological variation. *Nucleic Acids Res* 40: 4288–97.
